# Inference of Morphogen Gradient Precision from Molecular Noise Data

**DOI:** 10.1101/2021.04.20.440728

**Authors:** Roman Vetter, Dagmar Iber

**Affiliations:** Department of Biosystems Science and Engineering, ETH Zürich, Mattenstrasse 26, 4058 Basel, Switzerland; Swiss Institute of Bioinformatics, Mattenstrasse 26, 4058 Basel, Switzerland

**Keywords:** morphogen gradient, patterning precision, Sonic hedgehog, Bone morphogenetic protein, neural tube, progenitor number

## Abstract

During development, morphogen gradients provide spatial information for tissue patterning. Gradients and readout mechanisms are inevitably variable, yet the resulting patterns are strikingly precise. Measurement limitations currently preclude precise detection of morphogen gradients over long distances. Here, we develop a new formalism to estimate gradient precision along the entire patterning axis from measurements close to the source. Using numerical simulations, we infer gradient variability from measured molecular noise levels in morphogen production, decay, and diffusion. The predicted precision is much higher than previously measured—precise enough to allow even single gradients to define the central progenitor boundaries during neural tube development. Finally, we show that the patterning mechanism is optimized for precise progenitor cell numbers, rather than precise boundary positions, as the progenitor domain size is particularly robust to gradient alterations. We conclude that single gradients can yield the observed developmental precision, which provides new prospects for tissue engineering.

## 1 Introduction

During development, morphogen gradients convey positional information for tissue patterning. Measured morphogen gradients are typically approximated well by exponential functions,

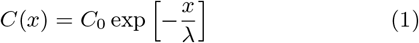

with amplitude *C*_0_ and decay length *λ* [3]. The length of the gradient, *λ*, specifies the distance over which the morphogen concentration decays by exp(1) ≈ 2.7-fold. The patterning domain is typically much larger than *λ* such that the concentration gradient declines substantially within. For instance, the Bicoid gradient length is a fifth of the length of early *Drosophila* embryos (*λ/L* = 0.2) [4]. The Decapentaplegic (Dpp) gradient in the *Drosophila* wing disc is even shorter (*λ/L* = 0.11) [5], and the relative lengths of the Sonic hedgehog (SHH) and Bone morphogenetic protein (BMP) gradients decline from about *λ/L* = 0.2 to *λ/L* = 0.05 during mouse neural tube (NT) development [2, 6]. In the developing vertebrate NT, positional information is provided by opposing SHH and BMP gradients [7] (Fig. 1A). Even if each morphogen patterns only one half of the domain, the morphogen concentration in the center of the final domain will be about 10^4^-fold lower than at the source. It is an open question whether morphogens can be sensed by cells over such distances, and how precise such patterning information would be.

**Figure 1:**
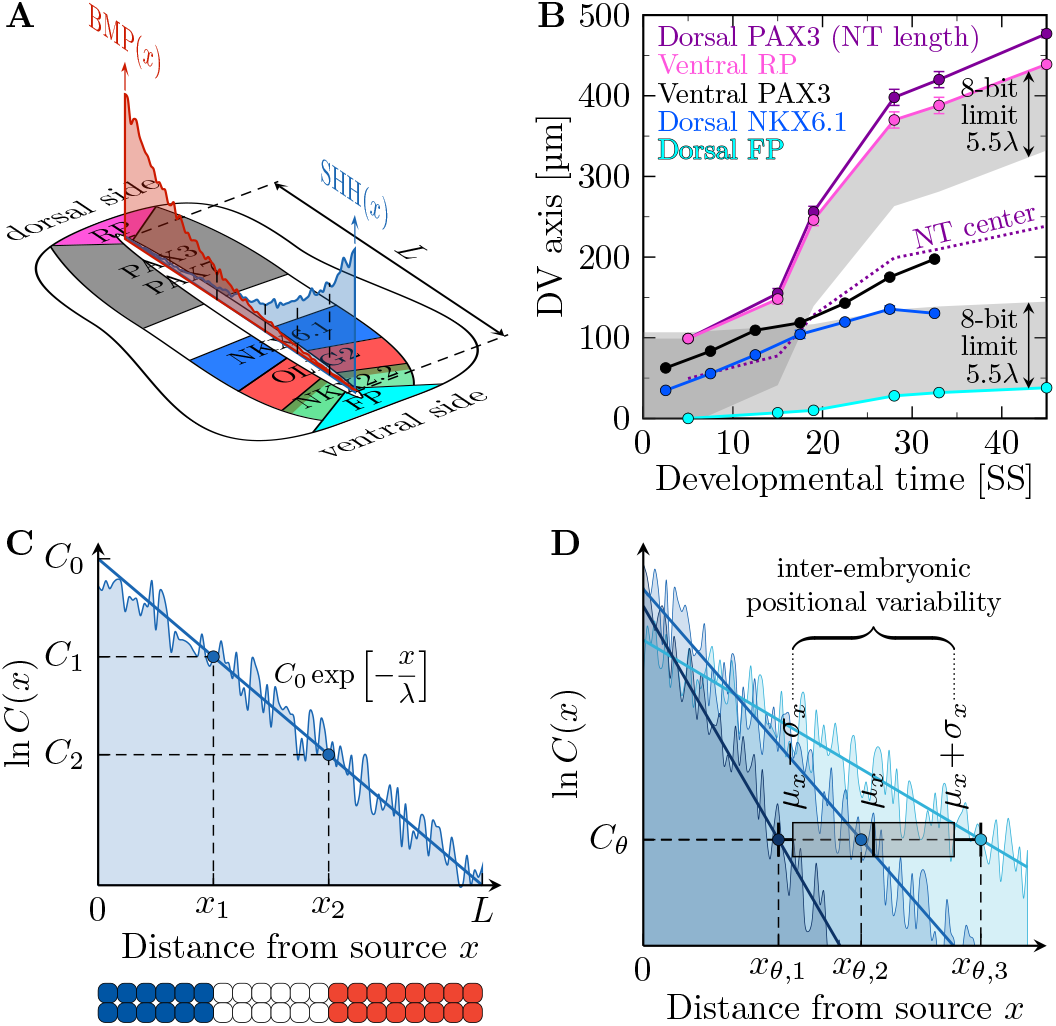
Positional error of gradients in the NT. **A** Schematic cross section of the mouse neural tube with noisy anti-parallel SHH and BMP gradients and emergent gene expression domains. The patterning domain of length *L* is confined ventrally by the floor plate (FP) and dorsally by the roof plate (RP). **B** NT length (purple) and position of domain limits over developmental time. At later stages, the dorsal NKX6.1 (blue) and ventral PAX3 (black) domain boundaries lie at the edge or outside the 5.5*λ* = 107 µm detection limit of 8-bit microscopy (shadowed). The pink and cyan lines mark the limit of the BMP-secreting RP and SHH-secreting FP respectively. The RP length was assumed to be equal to the FP length. Data points reproduced from [1, 2]. Error bars are standard errors. **C** According to the French flag model, the limits *x*_*θ*_ of distinct tissue domains are defined by concentration thresholds *C*_*θ*_. **D** Illustration of gradient variability between embryos. Different amplitudes and decay lengths yield different gradients (shades of blue), which translates to different positions *x*_*θ,i*_ where they attain a threshold *C*_*θ*_. This results in a distribution of readout positions with inter-embryonic mean *µ*_*x*_ and a positional error given by the standard deviation *σ*_*x*_.

In the bacterial chemotaxis response, adaptation allows cells to sense concentration gradients spanning at least five orders of magnitude, and cooperativity in receptor clusters enables a high gain such that the occupancy of one or two receptors can be sensed [8]. Whether similar effects are at work also in the NT is unclear, but adaptation in the SHH responsiveness has been noted [6, 9], and the PTCH1 receptor organizes as dimer of dimers, and each dimer binds one SHH [10]. Accordingly, it is, in principle, possible that morphogens can be sensed by cells over a 10^5^-fold concentration range (11.5*λ*), which corresponds to about 220 µm in the mouse NT—enough to cover the entire NT domain with opposing gradients (Fig. 1B). But even if the gradients can be sensed over 11.5*λ*, how precise would the conveyed positional information be?

According to the French flag model [11], the readout position *x*_*θ*_ is defined by a concentration threshold *C*_*θ*_ (Fig. 1C). By rearranging Eq. 1, one obtains

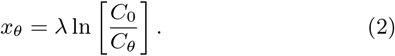

As a result of molecular noise, the gradient length *λ* and the ratio between gradient amplitude and readout threshold, *C*_0_*/C*_*θ*_, differ between embryos [2, 6, 12]. This difference translates into shifts in the readout positions *x*_*θ*_ (Fig. 1D). The overall readout position is typically calculated as the mean of the individual positions *x*_*θ,i*_ in the different embryos *i*, and the positional error is typically quantified by the standard deviation,

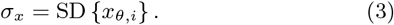

Following this definition, the readout position and accuracy of two centrally located progenitor domain boundaries have been quantified in the mouse NT [2]. The positional error of both the dorsal NKX6.1 boundary and the ventral PAX3 boundary was found to be about 1–3 cell diameters. In parallel, also the gradient variability was measured. Given the challenges in measuring morphogen gradients directly [6, 13], GBS-GFP was used as transcriptional reporter of SHH signaling, and phosphorylated Smad1/3/5 (pSMAD) as a readout of BMP signaling. Close to the source, the positional error increased from a single cell diameter (4.9 µm [6]) to about 3 cell diameters over time. In the center of the neural tube, however, the positional error was reported to increase from 3 cell diameters in early stages to more than 30 cell diameters later on. Combined readout of the imprecise SHH and BMP gradients was proposed to yield the higher precision of the central progenitor boundaries.

However, even a combined readout fails after 15 somite stages (SS), i.e., after about 30 hours of spinal cord development, and it remains unclear how the precise patterning of the central progenitor domains is achieved.

The detection of morphogens poses a challenge not only to cellular tissues, but also to microscopes. Standard protocols recommend the use of 8-bit images [14], which limits the maximal detection range to 2^8^ = 256-fold. For exponential gradients, this corresponds to an upper bound of about 5.5*λ*, which is 107 µm ≈ 22 cell diameters in the mouse NT (Fig. 1B, shaded region). In practice, the usable range will even be shorter, if technical noise occupies a few percent of the 8-bit channel. As the same settings are used in all measurements, the decline of the GBS-GFP and pSMAD gradient amplitudes over developmental time [2] further restricts the detection range at later times such that also the dorsal NKX6.1 boundary will lie outside the GBS-GFP detection range. While 16-bit imaging would be possible, at least the GBS-GFP reporter poses further limits. The GLI binding site that drives GBS-GFP expression stems from the *FoxA2* enhancer [15]. *FoxA2* is restricted to the ventral-most part of the neural tube (FP and p3 domain) because it requires very high SHH levels and depends on the activator form of GLI proteins [16]. Other SHH-dependent genes are well known to be expressed dorsally of the dorsal-most limit of the *FoxA2* domain (Fig. 1A). Accordingly, failure to express *FoxA2* more dorsally does not indicate absence of SHH, but too low SHH levels to drive its expression. While *FoxA2* expression is controlled by further regulatory interactions that will not impact on GBS-GFP, GBS-GFP is unlikely to report very low levels of SHH signaling that may still be sufficient to drive SHH-sensitive genes.

These difficulties in imaging morphogen gradients call for an alternative way to estimate gradient precision from measured data. We develop a theoretical framework and combine it with a computational approach here. Based on the measured variabilities in morphogen production, turnover and diffusion, we infer the variability of morphogen gradients and thus their positional error as it results from molecular noise. The resulting positional accuracy is consistent with the observed precision of the readout boundaries in the mouse NT. The gradients are thus, in principle, sufficiently precise to yield the observed patterning precision. Furthermore, we show that the size of gene expression domains is independent of the activity and variability thereof when defined by the threshold-based readout of a single morphogen gradient. This results in a very robust mechanism to produce precise numbers of progenitor cells.

## 2 Results

### 2.1 Estimation of positional error from statistical gradient properties

Close to the source, the measured gradients can be approximated well by exponential functions [2], and the positional error of the fitted exponentials is similar to that of the raw gradients [17]. In the following, we show how the positional error can be calculated from the summary statistics of the exponential gradients rather than by evaluating the standard deviation of individual gradients. This then allows to predict the positional error of the gradients at a distance from the source based on the observed variability closer to the source, assuming that the exponential gradient shape is maintained. With this formalism we can then infer the maximal gradient variability that would be consistent with the observed readout precision in the mouse NT.

The reported *λ* values for SHH in the mouse NT (Fig. 2A) are consistent with a (truncated) normal distribution (Fig. 2B). We therefore consider *λ* as a Gaussian random variable with mean *µ*_*λ*_ and standard deviation *σ*_*λ*_. While the mean value remains roughly constant at about 20 µm over developmental time (Fig. 2A), the deviation from the mean, as measured by the coefficient of variation CV_*λ*_ = *σ*_*λ*_*/µ*_*λ*_, has been reported to drop as the NT grows (Fig. 2C). At a given point in time (i.e., at a given size of the NT), the available data (Fig. 2D) suggests that the amplitude *C*_0_ is log-normally distributed (Fig. 2E). While the mean amplitude *µ*_0_ increases over developmental time (Fig. 2D), the deviation from the mean, as measured by the coefficient of variation CV_0_ = *σ*_0_*/µ*_0_, reportedly drops as the NT grows (Fig. 2F). The inferred statistical parameters are summarized in Tab. 1. Similar data for the opposing BMP gradient is not available, but once it so becomes, our formalism is likely to apply analogously to BMP, as it also diffuses into the NT.

**Figure 2:**
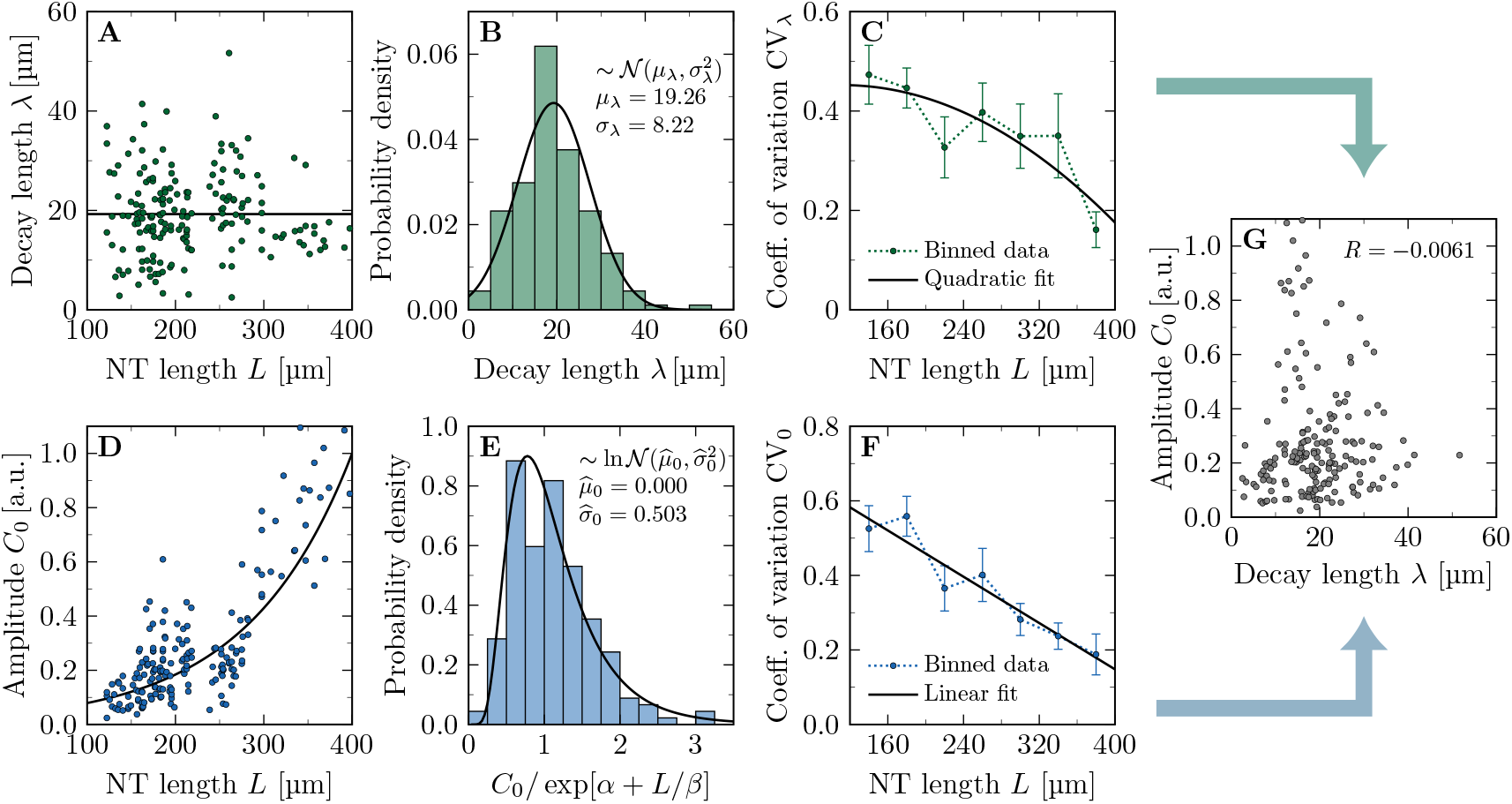
Statistical properties of the SHH gradient in the developing mouse neural tube. **A**,**B** *λ* is constant over developmental time, consistent with a truncated normal distribution. **C** Binning the data into 40 µm bins reveals that the relative variability in the data drops over time for SHH. **D** *C*_0_ increases as the neural tube expands. The solid line shows an exponential fit exp[*α* + *L/β*]. **E** Relative to the growing mean, the variability in the amplitude data is consistent with a log-normal distribution. **F** Also the relative amplitude variability of SHH declines over time. Error bars in **C**,**F** are bootstrapped standard errors. **G** SHH gradient length and amplitude are uncorrelated (Pearson correlation coefficient *R* ≈ 0). Data points reproduced from [6].

**Table 1:** Fitted statistical gradient parameters for the mouse neural tube.

| Gene/signal | Mean length $\mu_\lambda$ | Coeff. of variation $CV_\lambda = \sigma_\lambda/\mu_\lambda$ | Coeff. of variation $CV_0 = \sigma_0/\mu_0$ | Source |
| --- | --- | --- | --- | --- |
| SHH | 19.26 $\mu\text{m}$ | $-\left(\frac{L}{547.9 \mu\text{m}}\right)^2 + \frac{L}{1341 \mu\text{m}} + 0.410$ | $-\frac{L}{644.1 \mu\text{m}} + 0.769$ | [6] |
| GBS-GFP | 19.43 $\mu\text{m}$ | $-\left(\frac{L}{819.6 \mu\text{m}}\right)^2 + \frac{L}{861.0 \mu\text{m}} + 0.128$ | $\left(\frac{L}{790.8 \mu\text{m}}\right)^2 - \frac{L}{3437 \mu\text{m}} + 0.313$ | [2] |
| pSMAD | 22.67 $\mu\text{m}$ | $-\left(\frac{L}{820.3 \mu\text{m}}\right)^2 + \frac{L}{800.0 \mu\text{m}} + 0.072$ | $\left(\frac{L}{573.1 \mu\text{m}}\right)^2 - \frac{L}{722.2 \mu\text{m}} + 0.401$ | [2] |

Since morphogen concentrations are measurable only in arbitrary units (Fig. 2D) and since exponentials remain exponential independent of the chosen absolute scale, we can normalize the gradients by an arbitrary reference concentration without loss of generality. In the absence of precise knowledge about the readout threshold *C*_*θ*_, we choose the concentration scale such that *C*_*θ*_ = 1 in the following, which simplifies the notation. Our results retain their validity for general *C*_*θ*_. Hence, we assume that also the ratio *C*_0_*/C*_*θ*_ follows a log-normal distribution:

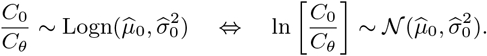

Here, 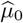 and 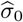 are the mean and standard deviation of the Gaussian random variable ln[*C*_0_*/C*_*θ*_]. We can use the properties of log-normal distributions [18] to express 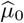 and 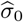 in terms of the mean *µ*_0_ and standard deviation *σ*_0_ of *C*_0_*/C*_*θ*_:

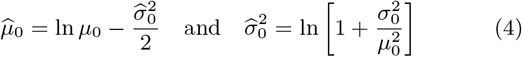

where

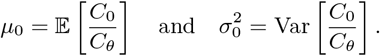

To estimate how domain boundaries behave under variability in the morphogen gradient, we seek to express the expected boundary position *µ*_*x*_ = 𝔼 [*x*_*θ*_] and its standard deviation *σ*_*x*_ = SD [*x*_*θ*_] as functions of the four gradient parameters *µ*_*λ*_, *σ*_*λ*_, *µ*_0_, *σ*_0_. The data for the SHH gradient in the mouse NT suggests that the gradient’s decay length and amplitude are uncorrelated (Pearson correlation coefficient *R* = –0.0061, Fig. 2G). This allows us to exploit the multiplicative properties of two independent random variables *X* and *Y*,

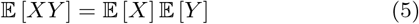

and

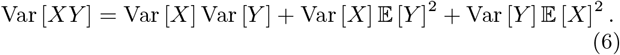

Putting Eqs. 2, 4 and 5 together, the mean boundary position is given by

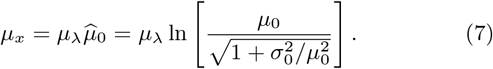

If there is no variability in *C*_0_*/C*_*θ*_ (i.e., 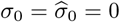), then Eq. 7 reduces to the deterministic case, Eq. 2.

The squared positional error follows from combining Eqs. 2, 4 and 6:

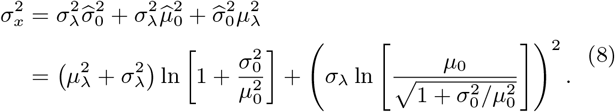

Notably, *σ*_0_ enters the position and positional error of a domain boundary only through the coefficient of variation CV_0_ = *σ*_0_*/µ*_0_. Eqs. 7 and 8 provide direct insight into how the statistical distributions of the gradient length and amplitude impact on the location and variability of the readout position. The larger *σ*_0_, the smaller *µ*_*x*_, i.e., the further the domain boundary shifts upward the concentration gradient, toward the morphogen source. Variability in the decay length *λ*, on the other hand, leaves the mean boundary position unaffected, as Eq. 7 is independent of *σ*_*λ*_. A larger mean gradient length or amplitude shifts the boundary downhill, away from the source. The positional error, on the other hand, depends on both gradient parameters and their scatter in a complicated nonlinear fashion that can even be non-monotonic. Tab. 2 and Fig. 3 summarize these relationships.

**Table 2:** Summary of the effect of concentration gradient parameters on domain boundary position and positional error.

| Parameter | Effect on $\mu_x$ | Effect on $\sigma_x$ |
| --- | --- | --- |
| $\mu_\lambda$ | positive | weak to positive |
| $\sigma_\lambda$ | none | positive |
| $\mu_0$ | positive | negative or weakly positive |
| $\sigma_0$ | weak to negative | weak to positive |

**Figure 3:**
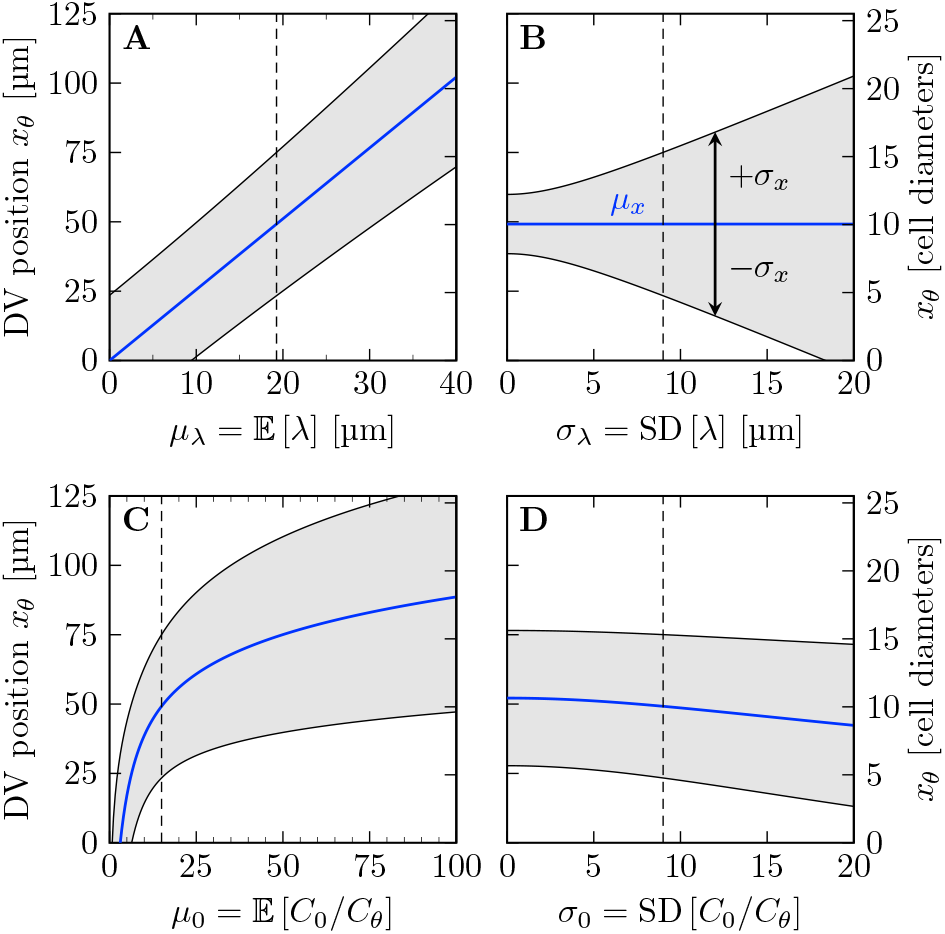
Effect of concentration gradient parameters on domain boundary position and positional error. Eqs. 7 (blue) and 8 (gray) are plotted. Each panel shows the variation of one parameter, with the other three fixed at measured early SHH values in mouse: *µ*_*λ*_ = 19.26 µm, *σ*_*λ*_ = 9 µm, *µ*_0_ = 15, *σ* _0_ = 9 (indicated by dashed lines).

We can further substitute Eq. 7 into Eq. 8 to obtain the positional error as an explicit function of the boundary position:

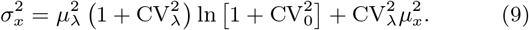

Eq. 9 is by construction identical to the direct way of computing the positional error via Eq. 3 [17] from infinitely many gradients.

It is worth noting that the positional error as a function of its readout position *µ*_*x*_, as given by Eq. 9, is independent of the mean gradient amplitude *µ*_0_. Precise knowledge of the change of *µ*_0_ over time (or as a function of *L*) is therefore not required to predict the positional accuracy in a noisy morphogen gradient. All that is needed is the variation of the amplitude relative to its mean, CV_0_. This has several beneficial consequences. A practical one is that no absolute measurement of the gradient amplitude is needed from experiments—relative values are sufficient to quantify positional accuracy. Another convenience is that the exact functional relationship used to fit or model the change of *µ*_0_ over time or length, be it exponential as in Fig. 2D, linear as in [6], or any other form, has no effect on the positional accuracy, as long as CV_0_ is given. Third, the fact that the absolute scale of the gradient amplitude is irrelevant implies that positional accuracy is unaffected by temporal changes in morphogen abundance, as long as CV_0_ remains sufficiently low.

### 2.2 Precision of gradient readout boundaries in the NT

With Eq. 9, the precision of a domain boundary is fully determined by its relative location in the patterning domain, *ξ* = *µ*_*x*_*/L*, the domain length *L*, the mean decay length *µ*_*λ*_, and the coefficients of variation of the gradient length and amplitude, CV_*λ*_ and CV_0_. As estimates for the latter three are known from measurements (Fig. 2, Tab. 1), we can predict the boundary precision in the growing NT at any point in development, anywhere in the patterning domain. For the reported gradient variabilities, the positional error in the center of the NT becomes as high as 15 cell diameters over time (Fig. 4A,B). The reported precision of the PAX3 and NKX6.1 domain boundaries (1–3 cell diameters) is more likely to be correct than that of the gradients as the steep boundaries and the concomitant change in the fluorescent signal are much easier to detect. We used Eq. 9 and numerical optimization to determine the variability at which the positional accuracy of the SSH and BMP gradients together, or one of them alone, would be consistent with the reported positional accuracy of the NKX6.1 and PAX3 domain boundaries. Assuming that the boundary position is always defined by the more precise gradient, we can reproduce the boundary precision with CV_*λ*_ = 0.08 ± 0.04, CV_0_ = 0.23 ± 0.33 for SHH and CV_*λ*_ = 0.06 ± 0.04, CV_0_ = 0.26 ± 0.16, 95% C.I. for BMP (Fig. 4C,E). Remarkably, fitting the reported boundary precision to the SHH gradient alone yields similar CV values, CV_*λ*_ = 0.05 ± 0.03, CV_0_ = 0.30 ± 0.20, 95% C.I. (Fig. 4D,E), challenging the previously proposed idea that opposing gradients serve to increase positional accuracy [2].

**Figure 4:**
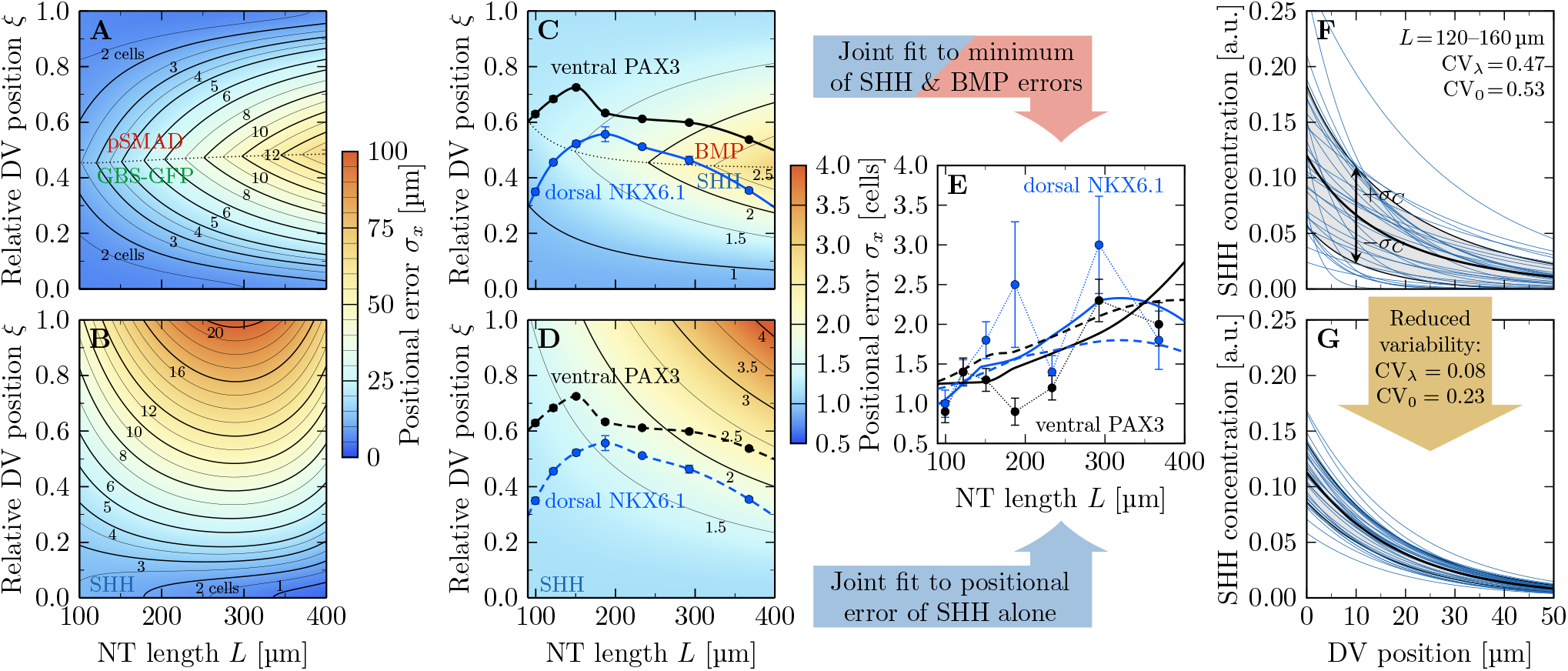
Gradient and readout imprecision in the developing neural tube. **A**,**B** With the reported gradient variability, the positional error of the opposing SHH and BMP gradients are in the order of several cell diameters. Eq. 9 is plotted using reported GBS-GFP and pSMAD (A) and SHH (B) parameters from Tab. 1. **C**,**D** Inferred positional error along the DV axis and over developmental time if the reported boundary precision in the dorsal NKX6.1 (blue) and ventral PAX3 (black) domain boundaries is matched, either by using always the more accurate of the SHH and BMP gradients (C), or if the SHH gradient alone determines the boundary positions (D). The dotted line in C divides the NT into two parts in which either SHH or BMP provide higher accuracy. Contour lines in A–D trace specified cell diameter values. **E** Reported (symbols) and predicted boundary precision if the domain boundaries are either set by SHH and BMP (solid lines, C) or by SHH alone (dashed lines, D). Domain boundary locations and errors in C–E are reproduced from [2]. Error bars were calculated as detailed in Methods. **F**,**G** The exponential SHH gradients with {*λ*_*i*_, *C*_0,*i*_} as reported in [2] are widely scattered in early NT development. The SHH gradients that match the measured positional error of the readouts (C,E) are still variable, but do not contain outliers. Shaded areas show standard deviations *σ*_*C*_. All error bars are standard errors.

The inferred CV values for the gradients lie near the lowest measured SHH gradient variability (Fig. 2C,F). Visual inspection shows that the reported variability corresponds to gradient profiles with some very short and some very long gradients that are difficult to reconcile with a successful patterning process (Fig. 4F). The variability inferred by us, while still resulting in highly variable gradients, does not result in such outliers (Fig. 4G). This raises the question whether the reported outliers reflect biological variation or technical problems in reliably measuring the morphogen gradients. Or differently put, how accurate are the reported gradient variabilities?

### 2.3 Technical limitations in measuring morphogen

According to the reported gradient properties (Tab. 1), the SHH gradient is only about half as precise as the GBS-GFP gradient (Fig. 5A), even though GBS-GFP is a direct SHH reporter [15]. We emphasize that this difference is observed already at the earliest developmental timepoint, long before adaptation results in the down-regulation of the SHH-dependent response [9]. Accordingly, this points to a higher technical variation for the SHH antibody staining than for the GBS-GFP reporter. The antibody staining for SHH in the NT patterning domain is very weak compared to that in the notochord, and gradient measurements are therefore challenging. Detection problems can be expected also with the GBS-GFP reporter as the *FoxA2* enhancer, that it is based on, responds only to the highest SHH concentrations [16]. In support of technical limitations in determining gradient variability, the coefficients of variation are strongly negatively correlated with the intensity of the signal for SHH and GBS-GFP (Fig. 5B,C), even though the gradient amplitude increases for SHH and decreases for GBS-GFP and pSMAD over developmental time due to adaptation [9, 19] (Fig. 5D), while the coefficients of variation show the opposite trend (Fig. 5E,F). This suggests that technical limitations at low concentrations artificially increase the reported variability, precluding an accurate measurement of the true gradient variability. We therefore turned to simulations to infer the expected variability based on the reported variability of morphogen production, degradation and transport rates.

**Figure 5:**
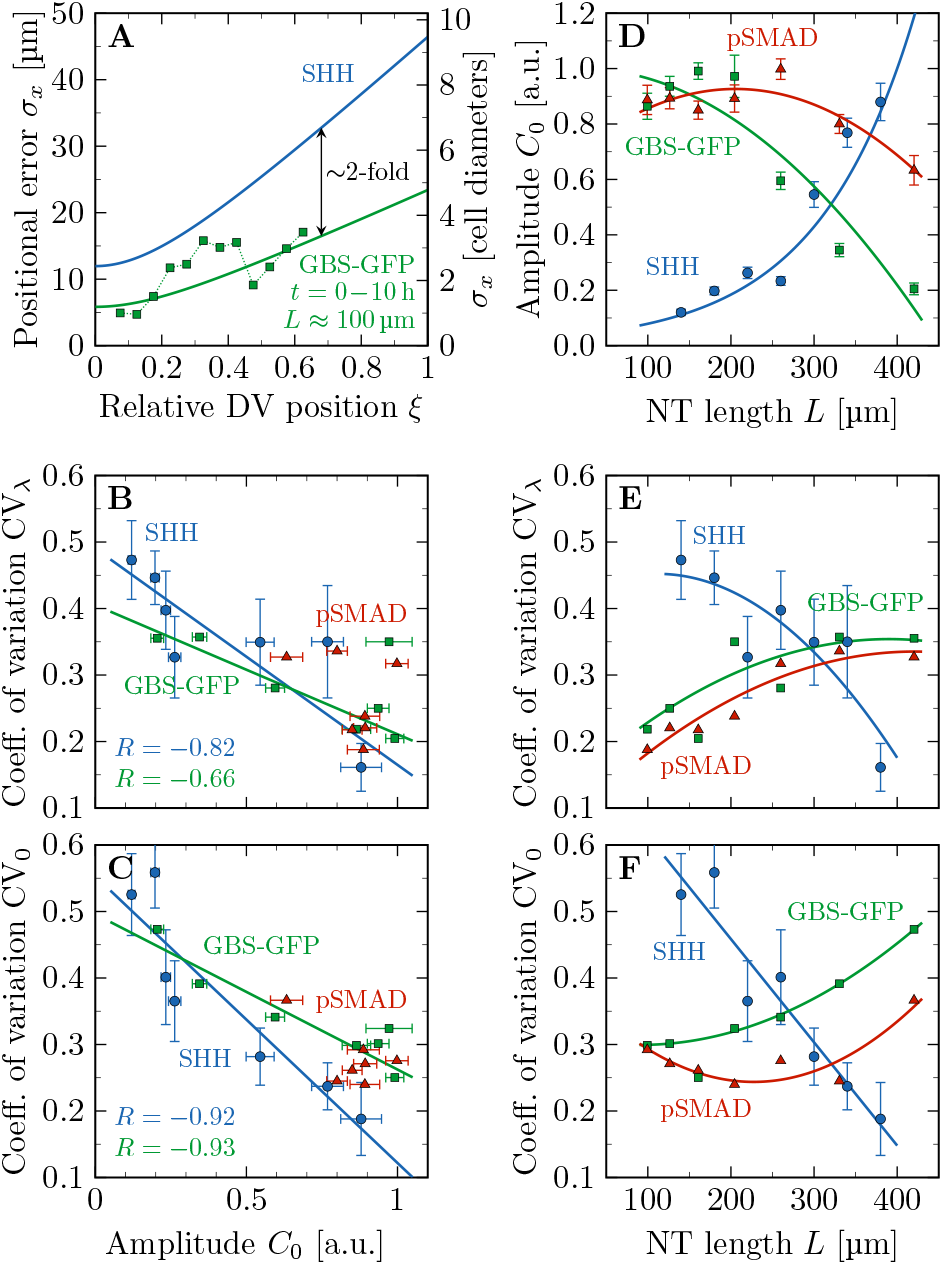
Technical limitations in measuring morphogen gradients. **A** With the reported variability, the SHH gradients would be about twice as imprecise as its readout GBS-GFP. The first developmental timepoint (0–10 h) of the GBS-GFP data (symbols) was reproduced from [2]; solid lines represent Eq. 9 with GBS-GFP and SHH parameters inferred from [2, 6] (Tab. 1). **B**,**C** Gradient variability is anti-correlated with the amplitude (Pearson correlation coefficient *R* ≪ 0), hinting at a potential technical limitation in the fluorescence intensity measurements. **D** The reported amplitudes of the SHH gradient increases, while the amplitudes of the GBS-GFP and pSMAD gradients decrease as the NT expands. **E**,**F** The reported gradient variabilities show the opposite trend. Solid lines are polynomial least-squares fits as listed in Tab. 1. Data in panels B–F was reproduced from [2, 6]; error bars are standard errors.

### 2.4 Gradient variability as a result of molecular noise

In a cellular tissue, the morphogen production, degradation, and transport rates vary from cell to cell. This variability ultimately generates the variability in the steady state morphogen gradient profiles. We can estimate this variability by simulating a simple reaction-diffusion model on a continuous 1D domain where these parameter values differ randomly from segment to segment (Fig. 6A). To describe the steady-state morphogen profiles, we solve the steady-state reaction-diffusion equation

**Figure 6:**
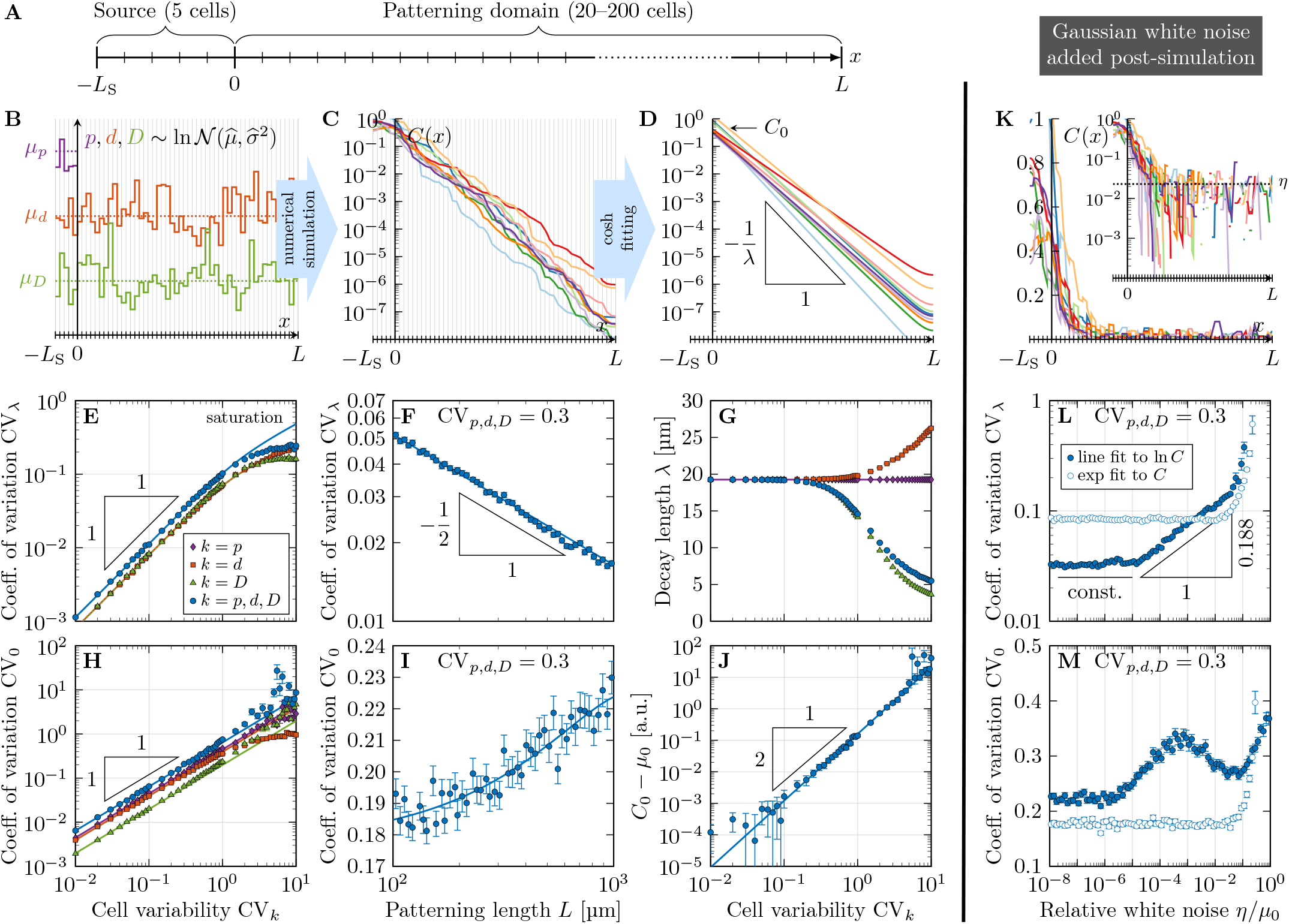
Numerical model predicts gradient variability from molecular noise. **A** Schematic of the simulated 1D domain. **B** Kinetic parameters *k* = *p, d, D* were drawn randomly and independently for each cell from log-normal distributions with specified mean *µ*_*k*_ and coefficient of variation CV_*k*_. **C** Solving the reaction-diffusion equation repeatedly yields noisy morphogen gradients, from which the decay length *λ* and amplitude *C*_0_ can be extracted by fitting hyperbolic cosines in the patterning domain (0 ≤ *x* ≤ *L*, **D**). **E** The resulting variability in *λ* grows linearly with the variability in the kinetic parameters as long as CV_*k*_ ≲ 1, and saturates as CV_*k*_ increases further. **F** The size of the domain length over which noisy gradients are fitted affects the variability of *λ* according to the law of large numbers, 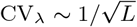 **G** Increasing molecular noise leads to a bias in the resulting fitted *λ*. **H** The amplitude variability also increases linearly with CV_*k*_, but does not saturate if all three parameters have a variability exceeding one. **I** The variability of the fitted amplitude moderately grows with increasing patterning domain length. **G** Noisy parameters also induce an overestimation of the amplitude deduced from fitting, proportional to 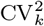. **K** Gaussian white noise ∼ 𝒩 (0, *η*^2^) added to the solution in all cells limits the range over which a line can be fitted to ln *C*. **L** Lin-fitting always (also at *η* = 0) leads to increased decay length variability, in particular with white noise stronger than one percent of the amplitude. Log-fitting is insusceptible to white noise as long as *η* ≲ 10^*−*5^*µ*_0_, and increases variability according to a power law with stronger white noise. If *η* exceeds a few percent of the amplitude, both fitting methods yield increased gradient length variability. **M** Amplitude variability is constant with lin-fitting for *η* ≲ 0.1*µ*_0_, whereas log-fitting yields larger CV_0_ values. *L* = 50 cells in all panels except F,I. All error bars are standard errors from 10^3^ independent simulations for each data point.

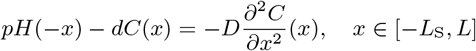

on a one-dimensional domain that was split into two subdomains, a morphogen source (−*L*_S_ ≤ *x* ≤ 0) and a patterning region (0 ≤ *x* ≤ *L*). The Heaviside step function *H* ensures that morphogen is produced at rate *p* only in the source, whereas it degrades at a linear rate *d* everywhere. Morphogen transport is driven by Fickian diffusion with diffusivity *D*. With zero-flux boundary conditions

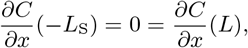

the deterministic solution is given by a concentration profile that follows hyperbolic cosines:

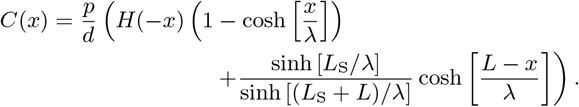

The decay length 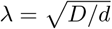 depends on the morphogen diffusivity *D* and the turnover rate *d*. The cosh is nearly exponential in the patterning domain except for a small deviation in the far end *x* ≈ *L* due to the zero-flux boundary. In the infinite size limit *L* → ∞, a pure exponential emerges for *x* ≥ 0:

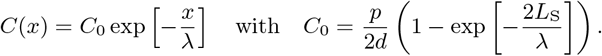

In our simulations, we divided both subdomains into cells of length 4.9 µm, the average cell diameter in the mouse NT [6] (Fig. 6A), and assigned each cell its own value of the three kinetic parameters *k* = *p, d, D*, drawn independently from log-normal distributions with prescribed means *µ*_*k*_ and coefficients of variation CV_*k*_ (Fig. 6B). Repeating the simulations many times for various CV_*k*_ values yielded independent noisy gradients spanning many orders of magnitude (Fig. 6C), from which we extracted *λ* and *C*_0_ by log-fitting hyperbolic cosines (Fig. 6D). We set 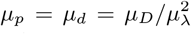 such that the deterministic decay length is the reported one of SHH, *µ*_*λ*_ = 19.26 µm [6]. The exact values of the parameters do not affect the steady state result as long as the relationship is maintained; we chose as mean parameter *µ*_*D*_ = 0.033 µm^2^/s as measured for Hedgehog (Hh) in the *Drosophila* wing disc [20] and fixed *µ*_*p*_, *µ*_*d*_ accordingly. The default setup consisted of 5 cells in the source, and 50 cells in the patterning domain.

This procedure yields the two gradient parameters and their variability as they result from molecular noise and NT expansion. We observe a linear increase of CV_*λ*_ as the cell variabilities CV_*d,D*_ are increased individually (keeping all others at zero), or all of them together, up to CV_*d,D*_ ≈ 1 (Fig. 6E). The production rate *p* affects only *C*_0_, not *λ*. This relationship can be understood theoretically. Since any product of powers of log-normal random variables is itself log-normal, so is 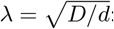:

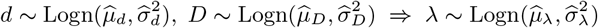

with

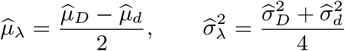

and

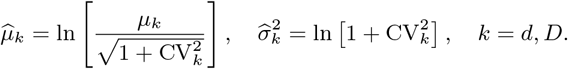

Using the expectation value and variance of log-normal distributions [18],

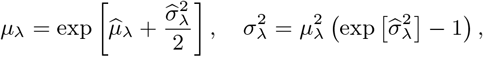

we find, for single cells,

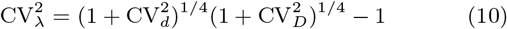

In patterning domains with many cells, CV_*λ*_ is lower due to cell averaging. The data from our simulations with *L* = 50 cell diameters precisely follows Eq. 10 up to CV_*d,D*_ ≈ 1, and in the case of *d* also beyond (Fig. 6E, lines), with a small proportionality constant shared by all curves. When all parameters are varied, though, CV_*λ*_ saturates at about 0.24. Larger values, such as the published CV_*λ*_ ≈ 0.4 for SHH (Fig. 2C), are unattainable even with extreme molecular variability, suggesting that the reported gradient variability [2, 6] is more technical than biological.

To examine the effect of the domain length *L*, we also varied the number of cells in the patterning domain from 20 to 200. As expected from the law of large numbers, log-fitting a variable exponential gradient over a longer domain leads to a more robustly fitted slope, such that 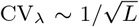 (Fig. 6F). This allows us to determine the size-dependent proportionality prefactor, resulting in

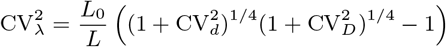

for CV_*D*_ ≲1, with fit parameter *L*_0_ = 6.13 ± 0.03 µm (mean ± SEM). Note that a declining CV_*λ*_ with increasing *L* is observed for the measured SHH gradient, but not for the GBS-GFP and pSMAD gradients (Fig. 5E), suggesting that amplitude effects perturbed the latter.

We further find that the fitted decay length starts to drift at moderate CV_*d,D*_ (Fig. 6G). If the morphogen diffusivity *D* or all parameters are noisy, *λ* is underestimated, whereas variability in the degradation rate *d* alone leads to overestimation of *λ*. This further attests to the difficulty in determining morphogen gradient parameters reliably from fitting noisy concentration profiles.

Unlike the decay length variability CV_*λ*_, the amplitude variability CV_0_ does not saturate as all cell variabilities are increased, but continues to grow linearly (Fig. 6H). We find CV_0_ to also grow mildly as the patterning domain lengthens (Fig. 6I). Finally, also the fitted amplitude is found to drift as molecular noise increases, proportionally to 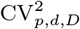 (Fig. 6J).

With these results, we can infer the physiological range of morphogen gradient variability by plugging in measured CV values. Quantitative data for the two morphogens are only available from measurements in the *Drosophila* wing disc. For Decapentaplegic (Dpp), the ortholog of mouse BMP4, CV_*d*_ = 0.5 has been reported for the degradation rate, CV_*p*_ = 0.59 for the production rate, and CV_*D*_ = 0.5 for the diffusion coefficient [21]. For Hh, quantitative data is available only for the diffusion coefficient, CV_*D*_ = 0.18 [20]. Measurements of other morphogens and in other species yield similar CV values [22, 23]. Single cell data is available only from cell cultures. The single-cell turnover rate variability of various proteins and transcription factors in mouse embryonic stem cells has been reported to be in the range CV_*d*_ = 0.16–0.45 [24]. For neural stem cell cultures only bulk measurements are available. Proteome half-life measurements yielded CV_*d*_ = 0.21 in mouse and 0.13–0.27 in human [25]. From protein half-life measurements in mouse neurons [26], one can infer a similar degradation rate variability of CV_*d*_ = 0.35–0.5.

Overall, the physiological range of inter-cell CV values appears to be 0.1–1, but most studies report CV *<* 0.6, and all these values likely include some technical noise. At an intermediate value of CV_*p,d,D*_ = 0.3, the biological gradient variability is CV_*λ*_ = 0.053, CV_0_ = 0.19 at *L* = 100 µm (CV_*λ*_ = 0.027, CV_0_ = 0.20 at *L* = 400 µm), which is precise enough to explain the NKX6.1 and PAX3 domain boundary errors by opposing SHH and BMP gradients, or even by SHH alone (cf. Fig. 4C–E). Even when we use a conservative CV value of 0.6 for all three kinetic parameters, the precision of a single morphogen gradient (CV_*λ*_ = 0.062, CV_0_ = 0.39) is consistent with the NKX6.1 and PAX3 domain boundary errors (1–3 cells).

We can further use the simulations to estimate the impact of technical limitations on the measured gradient variability. The measured gradients become noisy at about 5% of the maximal value [17]. We can represent this limitation by adding Gaussian white noise ∼ 𝒩 (0, *η*^2^) with uniform strength *η* to our simulated gradients in all cells, prior to fitting (Fig. 6K). The observed gradient variability strongly depends on the fitting method (Fig. 6L,M). Fitting the gradients in linear space (lin-fitting) always leads to elevated decay length variability, in particular for white noise exceeding 1%, but even at *η* = 0. Fitting the logarithmized gradients (log-fitting) yields significantly lower CV_*λ*_, but is insusceptible to white noise only as long as *η* ≲ 10^*−*5^*µ*_0_. At stronger white noise levels, we observe a power-law increase CV_*λ*_ ∼ *η*^*γ*^ with exponent *γ* = 0.188 ± 0.002 (SEM) and a cross-over with lin-fitting (Fig. 6L). If *η* exceeds a few percent of the amplitude, both methods yield significantly increased CV_*λ*_. Amplitude variability remains stable with lin-fitting for less than 10% white noise, whereas log-fitting yields mostly larger CV_0_ values (Fig. 6M).

In summary, our analysis suggests that natural noise in exponential morphogen gradients in the developing NT is sufficiently low to explain the previously reported progenitor domain boundary precision. Thus, both SHH and BMP gradients together— but even a single one of them alone—provide the spatial precision required to define the boundaries lying in the center of the NT with an error of only 1–3 cells. But can morphogen gradients provide even higher patterning accuracy for robust development?

### 2.5 Precision of progenitor domain size and progenitor number

In the vertebrate NT, the domain boundaries define the size of the different progenitor domains, which are formed as a result of different readout thresholds, as stipulated by the French flag model [11] (Fig. 7). Two domain boundaries located at *x*_1_ and *x*_2_ are the result of a morphogen readout at thresholds *C*_1_ = *C*(*x*_1_) and f_2_ = *C*(*x*_2_). As noted in [27], the length of a domain is given by

**Figure 7:**
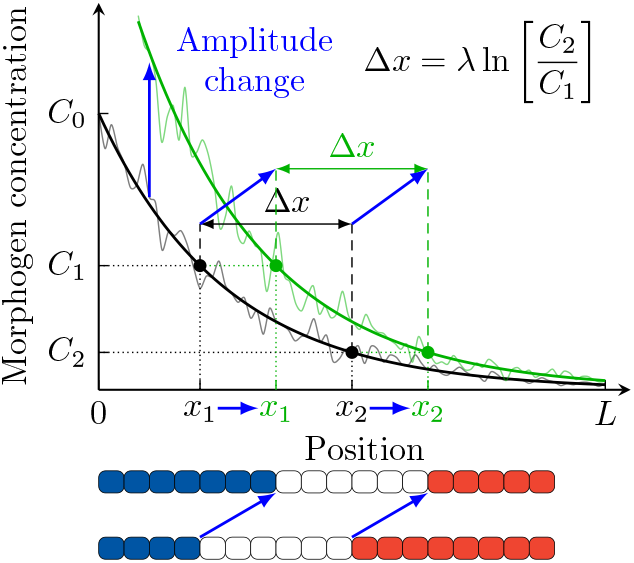
Robustness of patterning domain sizes to amplitude changes in the French flag model. The domain length Δ*x* = *x*_2_ − *x*_1_ is independent of the amplitude *C*_0_ of an exponential gradient. Amplitude changes therefore shift interior domain boundaries equally.

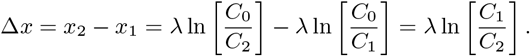

Notably, it is independent of the location in the entire patterning domain, and also independent of the gradient amplitude *C*_0_. Assuming that the domain width perpendicular to the *x* axis remains roughly constant along *x*, this paradigm provides a very robust mechanism to preserve the gene expression domain volume (and thus, the number of progenitor cells) during development. A change in the gradient amplitude *C*_0_ shifts both domain boundaries by the same distance, such that its size remains unchanged. The domain length is determined only by *λ*, which is stable over developmental time (Fig. 2A), and by the readout threshold ratio *C*_1_*/C*_2_. Only the very first and last progenitor domains in the pattern are affected by a transient amplitude *C*_0_, as one of their boundaries is given by the end points of the entire patterning domain, *x* = 0 and *x* = *L*.

Even in a probabilistic setting with variable gradients, the expected domain length *µ*_Δ*x*_ is unaffected by a change in amplitude, if *C*_1_, *C*_2_ and CV_0_ are constants:

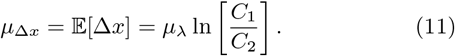

To quantify the variability of Δ*x* in a noisy gradient, we can calculate the variance

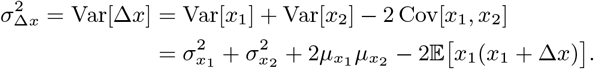

After some elementary algebra, assuming again independence of *λ* and *C*_0_ and using Eqs. 7, 8 and 11, all terms involving the amplitude cancel out, and we find the remarkably simple form

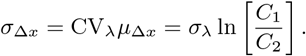

The inaccuracy of the size of a progenitor domain therefore scales with its size itself, with CV_Δ*x*_ = CV_*λ*_ as the proportionality constant. For an exemplary domain size of *µ*_Δ*x*_ = 50 µm and a coefficient of variation CV_*λ*_ ≈ 0.05, this results in a domain size error *σ*_Δ*x*_ as low as half a cell diameter, regardless of how far away from the source the domain lies. Strikingly, unlike their spatial boundary *positions*, the *length* of the gene expression domains is completely independent of variability in the gradient amplitude. The patterning mechanism thus appears optimized to generate precise progenitor cell numbers, rather than precise boundary locations. We emphasize that only a single morphogen gradient is required to achieve this high patterning precision.

## 3 Discussion

High patterning precision is pivotal for robust development. Given present technical limitations, the gradients can be reliably measured only within few *λ* from the source. We have now developed a formalism (Eq. 9) to estimate the positional error along the entire patterning axis from the gradient variability close to the source. We confirm previous reports that the experimentally determined gradient variability is too large to explain the high precision of the central domain boundaries in the mouse NT. Using computer simulations based on the reported variability in morphogen production, degradation, and diffusion, we find that the reported high variability of the gradient length *λ* cannot arise from natural noise in these parameters alone, as it saturates at lower values than previously measured for SHH, GBS-GFP and pSMAD, at high noise levels. With the gradient variability we inferred, the observed precision of the central progenitor boundaries is achieved even with a single gradient for the entire duration of the developmental process. Considering that the reported molecular noise levels are likely also elevated by technical errors, an even higher precision of the domain boundaries is still plausible. Finally, we show that the size of the morphogen-dependent tissue subdomains, that are not bordering the patterning domain edges, is even more precise than the individual boundary positions, because inaccuracies from amplitude variability cancel out. As a result, progenitors are produced in more accurate numbers than previously anticipated.

These insights provide novel perspectives on gradient-based patterning and on neural tube development in particular. Opposing morphogen gradients have been conjectured to be responsible for high patterning precision in a number of developing systems [2, 28, 29]. We now find that patterning in the NT can be controlled by a single gradient over the entire patterning period. Our simulations imply that the morphogen gradients remain exponential over a wide patterning distance. This will be the case if the same reactions and transport processes apply along the entire patterning axis. Measurements to confirm this are currently not available, and will require the development of more sensitive measurement technology. An important question is how cells can reliably detect very low morphogen concentrations, and what role adaptation plays in this [6, 8, 9].

Gradient variabilities have been reported also for other patterning systems, including Dpp and Wg in the *Drosophila* imaginal discs [5], and our formalism (Eq. 9) could thus be applied also in other developmental systems. We emphasize, however, that the accuracy of our formalism hinges on the accuracy of the measured gradient variabilities. Given how challenging it is to visualize morphogen gradients, technical errors are to be expected from such measurements. Based on the molecular noise simulations, we expect higher gradient precision also in other developmental systems.

Measuring the morphogen production, decay and transport rates is challenging, but is still easier than the detection of low morphogen concentrations, and thus offers a complementary approach to estimating gradient variability (Fig. 6). Current measurements of the morphogen production, decay and transport rates represent bulk measurements at the tissue level. Going forward, it will be valuable to obtain data on the single-cell variability of morphogen production and degradation rates. Currently, such data is available only from cell culture systems, but yields similar variabilities as bulk data. The reported CV values are in the range 0.1–1 across all species analyzed, including mice, flies, zebrafish, and humans. As this variability includes technical errors, these values present upper bounds. Even for a relatively pessimistic value of 0.6 for production, degradation and diffusion, we find that the gradient imprecision is 1–3 cells over several hundreds of micrometers, providing sufficiently accurate positional information to pattern a large domain. Local fluctuations can be reduced further through spatial and temporal averaging [30–32]. Moreover, in zebrafish, NT progenitor boundaries are sharpened by cell sorting [33, 34].

More than 50 years since the publication of the French flag model [11], it remains a matter of debate how morphogen gradients are read out [35]. Recent experiments support a threshold-based readout of the BMP gradient along the zebrafish dorsal-ventral axis [36]. Our finding that single noisy gradients provide a much more robust positional patterning mechanism than previously appreciated resolves the long-standing conundrum of how the observed patterning precision can be achieved with a simple threshold-based readout of a single gradient. This opens new avenues for tissue engineering, which simplifies substantially if single gradients suffice to do the job. The presented formalism is not limited to vertebrates or flies, but applies directly also to all other morphogen-dependent patterning systems in which morphogen transport is essentially diffusive.

## Methods

### Gradient data

The individual *λ*_*i*_ and *C*_0,*i*_ for SHH, as well as the sample means and standard deviations *µ*_*λ*_, *σ*_*λ*_, *µ*_0_, *σ*_0_ for GBS-GFP and pSMAD were extracted from the respective publications [2, 6].

### Numerical optimization of gradient variability

We determined the gradient parameter variabilities CV_*λ*_ and CV_0_ in Fig. 4 by fitting Eq. 9 to the progenitor domain boundary errors with MATLAB’s nonlinear least-squares curve fitting routine `lsqcurvefit`.

### Inference of error bars for the positional boundary error

In Fig. 4E, we inferred the uncertainties (error bars) associated with the positional error of the domain boundaries assuming that the boundary positions are normally distributed. In this case, the standard error of the standard deviation *σ*_*x*_ is given by 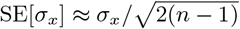 where *n* is the sample number [37].

### Simulation of gradient variability from molecular noise

The reaction-diffusion equation was solved with MATLAB’s boundary value problem solver bvp4c with absolute and relative error tolerances of 10^*−*10^. At each interface between two adjacent cells, continuity of the morphogen concentration *C* and its flux −*D∂C/∂x* was imposed.

## Code Availability

The complete source code for Fig. 6 is publicly released under the 3-clause BSD license. It is available as a git repository at https://git.bsse.ethz.ch/iber/Publications/2021_vetter_gradient_variability.

## Acknowledgements

We thank Marius Almanstötter, Marcelo Boareto, James Briscoe, Josée Dias, Johan Ericson, Thomas Horn, Alice Kluzer, and Marco Kokic for discussions.

## Author Contributions

RV & DI conceived the study, analyzed the data, and wrote the manuscript, RV developed the theory and simulations and produced the figures.

